# Loss of the Y chromosome is transcriptionally silent in clear cell renal cell carcinoma and across cancer entities

**DOI:** 10.64898/2026.10.08.757570

**Authors:** Alex Waterhölter, Rebecca Scholz, Andreas Tsouris, Francesca Cuomo, Mojca Adlesic, Léonel Nguekeu Zebaze, Christian Witt, Zulema Rodríguez Hernández, Anna Köttgen, Melanie Boerries, Pascal Schlosser, Ian J. Frew

## Abstract

**Background:** Loss of the Y chromosome (LOY) is the most frequent somatic alteration in male cancers and has recently been linked to immune evasion, metabolic reprogramming and prognosis. Because the Y chromosome encodes transcriptional and epigenetic regulators, such as UTY and KDM5D, LOY is assumed to act through dysregulation of autosomal gene expression. We asked whether LOY produces a transcriptional signature in clear cell renal cell carcinoma (ccRCC), where about 40% of male tumors lose the Y chromosome.

**Methods:** Here, we inferred Y chromosome status from RNA-sequencing of male ccRCC in TCGA-KIRC (n = 289) and CPTAC-ccRCC (n = 114), in 32 tumors from an autochthonous Vhl/Trp53/Rb1-deficient mouse model, and in five isogenic Y-deficient and five Y-retaining clones of the 786-O cell line. Differential expression was tested with Wilcoxon rank-sum tests on age- and purity-adjusted residuals combined by Stouffer meta-analysis (human cohorts) or with DESeq2 (mouse, cell clones), followed by gene set enrichment analysis. Pseudoautosomal genes, present on both sex chromosomes, served as internal controls for a halving of gene dosage.

**Results:** In all three settings, transcriptional changes were confined to the expected dosage effects of male-specific Y and pseudoautosomal genes. No autosomal gene reached the significance level of the non-compensated pseudoautosomal genes in human ccRCC, 14 autosomal genes were altered in the mouse model and 40 in the isogenic 786-O clones, none shared across settings. Gene set enrichment analysis revealed a modest, reproducible upregulation of cell division, ribosome biogenesis and DNA repair pathways shared between human and murine ccRCC. A meta-analysis across 27 male TCGA cancer entities (1,710 LOY, 2,988 Y-retaining tumors) recovered the same pattern with no autosomal gene showing a consistent large effect, and the same cell division, ribosome biogenesis and DNA repair pathways were modestly upregulated. Cancer/testis antigen expression was elevated in LOY tumors of several entities, most notably stomach adenocarcinoma, but not ccRCC.

**Conclusions:** Beyond direct gene dosage, LOY leaves the transcriptome of ccRCC, and of most other cancer entities, essentially unchanged. Y-encoded genes are therefore unlikely to drive cancer through direct transcriptional regulation of autosomal genes, directing attention to non-transcriptional effects of LOY.

## BACKGROUND

The Y chromosome is one of the smallest human chromosomes. Its approximately 62 million base pairs encode only 106 protein-coding genes [1], many of which are essential for male sex determination and reproductive function. For more than half a century, it has been known that blood cells tend to lose the Y chromosome as men age, termed mosaic loss of Y (mLOY) [2, 3]. This phenomenon was for a long time regarded as a benign consequence of aging. More recent studies, however, have revealed that mLOY in blood is associated with increased susceptibility to a diverse set of age-related pathologies, including cancer [4], cardiac fibrosis, heart failure [5], and Alzheimer’s disease [6]. Why Y chromosome loss is associated with pathologies across such diverse tissues remains unclear. The combined loss of several Y-chromosome-encoded proteins, including the transcriptional and epigenetic regulators UTY, KDM5D and ZFY, is believed to cause transcriptional dysregulation that might contribute to disease pathologies [7]. In keeping with the idea that certain Y chromosome-encoded genes carry out essential functions, it is notable that several, such as UTY, KDM5D and DDX3Y, have functional homologs that escape X inactivation on the X chromosome [8]. This argues that the dosage of these genes matters, and that in male cells it is ensured jointly by the X and Y homologs.

In addition to mLOY in blood cells, loss of the Y chromosome (LOY) is the most frequent genetic alteration in both hematologic and solid tumors in men [9, 10]. Recently, LOY was proposed to represent an immune-escape mechanism in bladder cancer, providing the first functional mechanism for a causal role of LOY in solid tumor biology [11]. LOY carries prognostic value in selected cancer types such as uveal melanoma, while in others it does not [12]. Additionally, a link between LOY in immune cells and cancer has been established: LOY in both tumor cells and T cells is associated with a more aggressive phenotype across epithelial cancers [13]. The underlying mechanisms driving these associations and their generalizability across tumor types and cell lineages remain unclear [14].

Clear cell renal cell carcinoma (ccRCC) is the most common adult renal malignancy with approximately 40% of male cases exhibiting LOY [12]. Although LOY is a frequent event in ccRCC, it remains unclear whether it contributes to disease biology. We therefore asked whether LOY produces a distinct transcriptional signature in ccRCC. To address this, we analyzed human and mouse ccRCC tumors and generated isogenic pairs of Y-retaining and Y-deficient ccRCC clones.

## METHODS

### Cell culture

786-O cells (RRID:CVCL_1051, male) were obtained from ATCC. Cells were cultured in RPMI-1640 (Gibco) supplemented with 10% FCS (Anprotec) and 1% Penicillin/Streptomycin (Gibco). MuLE vectors (RRID:Addgene_62178) expressing sgRNAs, spCas9 and NeoR were cloned and packaged into lentiviruses that were used to infect cells as previously described in [15]. Transduced cells were selected using neomycin (G418 disulfate salt, 500 µg/mL, Sigma-Aldrich, G8168) within 10 days. The three control sgRNA sequences (5’ - 3’) were sgCtrl (GTCATGTCACTTATCAAGTC), a non-targeting scramble RNA control, sgRBMY1A1 (GCATATAGTAATACACGAGA), targeting the ampliconic Y chromosomal gene RBMY1A1 and sgY-Cen (CGGGAGTTCAACCTATCATT) targeting alpha-satellite DNA in the Y centromere. sgRBMY1A1 and sgY-Cen were designed to lack an adjacent PAM motif and represent binding but non-active control sgRNAs. sgTSPY (GTATCCGGATTATGAAGTGG), targeting the ampliconic Y chromosomal gene TSPY, represents an active sgRNA. 786-O WT cells used for cloning were tested negative for mycoplasma most recently on 10/17/24 with a qPCR-based method (Eurofins). 786-O-WT cells were confirmed as 786-O by STR (Eurofins Human Cell Line Authentication Service-STR DNA profiling). Generated clones were passaged a maximum of 35 times.

### Generation of single cell clones

Single cells were seeded in 96-well plates and cultured in 786-O wild-type-conditioned RPMI-1640 for ∼14 days. For expansion of the clonal populations normal RPMI-1640 was used.

### Western Blot

For sample preparation, cells were washed once with ice-cold PBS, scraped from culture plates in PBS, pelleted and lysed in RIPA buffer supplemented with NaF, Na_3_VO_4_ and Protease inhibitor cocktail (Sigma-Aldrich). Protein concentrations were determined using BCA assay (ThermoFisher). SDS-PAGE was performed under standard conditions. Proteins were transferred to a nitrocellulose membrane (Cytiva) in a semi-dry blotting system (Scie-plas) using cold tris-glycine transfer buffer with 20% methanol. Membranes were blocked with TBS with 5% non-fat dry milk (ITW Reagents) for 1 hour at room temperature and then incubated with the primary antibody. Antibody solutions were prepared in TBS-Tween (0.1%) with 5% non-fat dry milk and incubated overnight at 4°C (primary antibodies) or for 1 hour at room temperature (anti-Vinculin and secondary antibodies). Membranes were washed three times for 5 min with TBS-T after each antibody incubation. Odyssey DLx Imaging System was used for antibody detection. The following antibodies were used: anti-KDM5D (Bethyl Laboratories, A301-751A, 1:1000, RRID:AB_1211352), anti-Cas9 (Cell Signaling, #14697, 1:1000, RRID:AB_2750916), anti-Vinculin (abcam, ab129002, 1:1000, RRID:AB_11144129), goat anti-mouse IgG IRDye® 800CW (LICORbio, 926-32210, 1:10000, RRID:AB_621842), and goat anti-rabbit IgG IRDye® 800CW (LICORbio, 926-32211, 1:10000, RRID:AB_621843).

### RT-qPCR

Total RNA was extracted from cells using the NucleoSpin RNA Kit (Macherey-Nagel). First-strand cDNA synthesis was done using 2 µg of RNA with the Ready-To-Go You-Prime First-Strand Beads (Cytiva). Quantitative PCR was performed in technical triplicates using Takyon SYBR 2X MasterMix blue dTTP with fluorescein and a LightCycler 480 System (Roche). Primer sequences (5’ – 3’) were *DDX3Y*_for (CGGCAGTAACTGTCCTCCACAT), *DDX3Y*_rev (CTGGAGTAGGACGAGTATAGCG), *KDM5D*_for (CGCAGCTTTGAAGAGCTAAG), *KDM5D*_rev (CAGCTGTGGAGTGTCCATCC), *RPS4Y1*_for (TCTTCCGTCGCAGAGTTTCG), *RPS4Y1*_rev (TGAGGAAGACGATCAGAGGAA), *RPS18*_for (GCGGCGGAAAATAGCCTTTG), *RPS18*_rev (GATCACACGTTCCACCTCATC). Relative expression normalized over *RPS18* expression and an internal control was calculated using the Livak method (2^-ΔΔCt^).

### RNA-seq from cell lines

RNA was isolated from five selected distinct ROY and LOY 786-O clones each (biological replicates) from cells grown to 70% confluency using the NucleoSpin RNA Mini kit (Macherey-Nagel). Quality control, poly-A enrichment and sequencing of mRNA libraries on a NovaSeq X Plus platform (Illumina) using paired-end 150 bp sequencing were conducted by Novogene GmbH in Munich, Germany.

### Human cohort resources

Male-derived tumor samples were drawn for three public cohorts: TCGA-KIRC (n = 338), CPTAC-ccRCC (n = 137) and TCGA pan-cancer. For both ccRCC cohorts, raw gene expression counts and clinical metadata were obtained from the GDC harmonized pipeline (https://gdc.cancer.gov/). For TCGA pan-cancer data, expression was obtained from the TCGA pan-cancer Atlas EBPlusPlusAdjustPANCAN_IlluminaHiSeq_RNASeqV2.geneExp.tsv resource (https://gdc.cancer.gov/about-data/publications/pancanatlas) [16]. TCGA-BLCA (n = 224) was drawn out of the pan-cancer data. Somatic mutation calls were obtained from cohort-matched WGS/ WES-der ived MAFs f or t he ccRCC cohor t s and f r om t he MC 3 public MAF (mc3.v0.2.8.PUBLIC.maf.gz) [17] for BLCA. Expression-based tumor purity estimates (ESTIMATE) were taken from the TCGAbiolinks (v2.38.0, RRID:SCR_017683) [18] Tumor.purity resource for the TCGA cohorts and from Li et al. [19] for CPTAC. A cancer/testis gene list was obtained from Wang et al. [20], Supplementary Data 3A (1,019 genes, of which 954 were quantified in the TCGA pan-cancer matrix).

### Human cohort RNA-seq analysis

For differential expression analysis in TCGA KIRC and CPTAC ccRCC, per-cohort counts were filtered to genes with at least 10 reads in at least 15% of samples. For BLCA, we retained genes with a log2(RSEM+1) > 1 in at least 15% of the samples and restricted to protein-coding genes. Pan-cancer cohort analysis was restricted to protein-coding genes which are testable in 90% of all cohorts (16,919 genes). We size-normalized the TCGA and CPTAC ccRCC counts using DESeq2 (v1.50.2, RRID:SCR_015687) [21]. Then, we log2-transformed and regressed out age and tumor purity, which has been shown to confound interpretation of RNA expression [22] (residuals from y ∼ age + tumor_purity, re-centered on the gene mean). A two-sided Wilcoxon rank-sum test was applied to the residuals between LOY and ROY (retention of Y chromosome) groups within KDM5C-wild-type (for ccRCC) or *KDM6A*-wild-type (for BLCA) samples, and per-gene log2-fold changes were computed as the difference of group means. No mutation filtering was performed on the pan-cancer cohort level analysis. We applied this non-parametric approach since human cancers are subject to numerous copy-number alterations that violate the distributional assumptions of parametric differential expression algorithms and thus have been shown to produce increased false-positives [23]. For ccRCC, per-cohort statistics were combined across TCGA-KIRC and CPTAC-ccRCC using a sample-size-weighted Stouffer’s method:

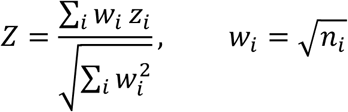

where

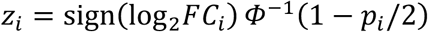

is the signed Z score of cohort i, *n_i_* is the number of tumors analyzed in cohort i, and *Φ*^−1^ is the inverse standard normal cumulative distribution function. Fold change is reported as the sample-size-weighted mean:

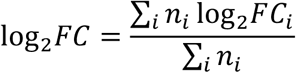

Analyses were restricted to the 16,178 genes quantified in both cohorts (protein-coding genes plus XIST). Significance is reported as the Benjamini–Hochberg false discovery rate (FDR). For the pan-cancer analysis LOY status was taken directly from WGS/WES-based Qi et al. [12], with LOY defined as the union of LOY, partial-LOY and relative-LOY calls and ROY as Y-intact (N = 1,710 LOY, N = 2,988 ROY). The same above mentioned non-parametric procedure was applied independently within each of the 27 male TCGA entities. Adjustment used the purity and age covariates only when available. Tumor purity estimates are absent for 11 of the 27 entities (CHOL, DLBC, ESCA, LAML, MESO, PAAD, PCPG, SARC, STAD, TGCT and UVM), in which we therefore regressed out only age. Per-entity statistics were then combined by the same sample-size-weighted Stouffer’s Z across the entities. Significance is reported as Benjamini-Hochberg FDR < 0.05.

### Mouse RNA-seq analysis

Publicly available raw RNA-seq reads (GSE344622, GSE330887, GSE150983, [TBD]) of VpR mouse ccRCCs from our previous experiments were retrieved and trimmed with *trimmomatic* (v0.39, RRID:SCR_011848) [24] with the options *LEADING:3 TRAILING:3 SLIDINGWINDOW:4:15* to remove bases with very low quality and *MINLEN:25* to remove small read fragments. We aligned the trimmed reads to the GRCm39 reference genome using *STAR aligner* (v.2.7.10a) [25] with the following parameters *--outFilterMultimapNmax 100 --alignSJoverhangMin 8 --alignSJDBoverhangMin 1 --out Filter MismatchNmax 999 --out Filter MismatchNover Lmax 0. 05 --alignIntronMin 20 --alignIntronMax 1000000 --alignMatesGapMax 1000000 --outFilterMatchNmin 16*. Read quality was assessed with FastQC (RRID:SCR_014583) before and after trimming. Gene expression was then quantified with the *featureCounts* command of *Rsubread* (v2.24.0, RRID:SCR_009803) [26] and the genes of the GRCm39 (Ensembl annotation) reference genome. We then inferred the presence of the Y chromosome by calculating a Y-score as the mean of the gene-wise z-scores of Y chromosome genes having a minimum expression level of 10 counts in the healthy samples. The lowest Y-score of the healthy samples was selected as the threshold for LOY and the highest normal murine kidney cortex samples’ Y-score as an upper limit to eliminate any samples with potential duplication or large structural variants in the Y chromosome. We eliminated genes with count values under 10 in less than 15% of the samples and performed Principal Component Analysis (PCA) using prcomp and the transcript per million (tpm) expression levels excluding genes of the Y chromosome for the PCA. To assess the differential expression of genes between LOY and ROY samples we used *DESeq2* (v1.50.2, RRID:SCR_015687) [21] and genes with a total count less than 1000 were removed.

### 786-O RNA-seq analysis

Raw RNA-seq reads were initially cleaned with *trimmomatic* (v0.39) and then aligned to the GRCh38 reference genome using *STAR aligner* (v.2.7.10a, RRID:SCR_004463) [25] with the same parameters as in the Mouse RNA-seq analyses section. Read quality was assessed with FastQC (RRID:SCR_014583) before and after trimming. Gene expression was then quantified with the *featureCounts* (RRID:SCR_012919) command of *Rsubread* (v2.24.0, RRID:SCR_009803) [26] and the genes of the GRCh38 (Ensembl annotation release 110) reference genome. We allowed multimapping reads with the *featureCounts* parameters *allowMultiMappingReads = TRUE* and *allowMultiOverlap = TRUE* in order to capture the expression of PAR genes that are present both on the X and Y chromosomes. We eliminated genes with count values under 10 in more than 50% of the samples and performed Principal Component Analysis (PCA) using prcomp and the transcript per million (tpm) expression levels. To verify the loss of the Y chromosome we computed a Y-score as the median of z-scores of all male-specific Y chromosome genes that have a minimum count of 10. Differential expression between LOY and ROY samples was assessed with DESeq2 (v1.50.2, RRID:SCR_015687) [21] and genes with a total count less than 1000 were removed.

### Gene-set enrichment analysis

fgsea (v1.36.2, RRID:SCR_020938) [27] was run on the meta rank metric sign(log2FC) × −log10(max(p, 1e−300)) against Gene Ontology gene sets (biological process, molecular function and cellular component) retrieved from GO.db (v3.22.0, RRID:SCR_024241)[28], gene sets were restricted to 10-500 members, MSY genes were removed from every set and sets falling below 10 members were discarded (nPermSimple = 10000, seed 42). Gene mapping was achieved through org.Hs.eg.db (v3.22.0, RRID:SCR_024739) [29] for human and org.Mm.eg.db (v3.22.0, RRID:SCR_023488) [30] for mouse gene sets. Bonferroni adjustment was applied within each cohort’s term universe. Terms with a Bonferroni-adjusted-p-value < 0.05 were considered significant.

### Grouping of enriched GO terms into functional themes

Bonferroni-significant GO terms from each GSEA were grouped into higher-level themes by the overlap of their leading-edge gene sets. For every pair of significant terms, we computed the Jaccard index of their leading-edge gene lists, J(A,B) = |A ∩ B| / |A ∪ B|. Terms were then clustered by average-linkage hierarchical clustering on the distance 1 − J and the dendrogram cut at a height of 0.85, so that terms grouped into a theme share a leading-edge Jaccard index of at least 0.15. Themes were visualized as treemaps in which tile area is proportional to the number of genes annotated to each term within the ranked list and color denotes theme membership. Visualization was performed using the R package treemapify (v2.6.0).

### DepMap dependency analysis

CRISPR gene-effect (Chronos) scores, RNA expression (log2(TPM + 1)) and sample annotations were obtained from DepMap release 25Q2. Male ccRCC (n = 9) and male bladder urothelial carcinoma (n = 17) cell lines were classified as LOY or ROY using median z-scored expression of the same nine MSY scoring genes as in the human cohort analysis. Within each panel, testing was restricted to the X-encoded gametologs of these MSY genes that were expressed in that panel (log2(TPM + 1) ≥ 1 in at least 50% of lines), yielding 11 gametologs (*DDX3X, EIF1AX, KDM5C, KDM6A, RPS4X, TMSB4X, USP9X, ZFX, PRKX, TBL1X* and *TXLNG*) in the ccRCC panel. Gene-effect scores were compared between LOY and ROY lines with Welch’s two-sided t-test and Benjamini-Hochberg FDR correction across the tested gametologs.

### Statistical Analysis

Sample sizes were not set by a power calculation. We therefore assessed sensitivity *post hoc*. Dispersions were recovered from the unshrunken standard errors reported by DESeq2 for expressed genes (baseMean > 100) using Var(log FC) ≈ (1/n₁ + 1/n_2_)(1/μ + φ), giving φ = 0.178 for the murine VpR tumors (14 LOY, 18 ROY) and φ = 0.033 for the isogenic 786-O clones (5 LOY, 5 ROY). Natural-log values were converted to log_2_ throughout. At 80% power against the realized Benjamini-Hochberg thresholds (0.05 FDR; p < 7.0 × 10^-5^ and 2.4 × 10^-4^) and at median expression, we had power to resolve an absolute log₂ fold change > 1.05 in the murine cohort and > 0.77 in the cell line. For the human cohorts, analyzed by Wilcoxon rank-sum test on covariate-adjusted residuals and combined by sample-size-weighted Stouffer’s Z, the corresponding sensitivity is a standardized effect of 0.44 SD at the realized Benjamini-Hochberg threshold (p < 1.3 × 10^-3^). Genes at the significance boundary have a median absolute log₂ fold change of 0.17, indicating that the human ccRCC meta-analysis resolves autosomal effects of approximately 12% or greater. Samples were not randomly assigned to groups, as group membership (LOY versus ROY) is an intrinsic property of each tumor or clone rather than an assigned treatment. Investigators were not blinded to Y chromosome status. For comparisons with fewer than five samples per group, individual data points are plotted.

### Software

Murine heatmaps were generated using the R package ComplexHeatmap (v2.26.1, RRID:SCR_017270) [31] while all other visualizations, when not stated differently, were generated using ggplot2 (v4.0.2, RRID:SCR_014601) with ggrepel (v0.9.8) for label placement, patchwork (v1.3.2) for panel composition, and extrafont (v0.20) and showtext (v0.9-8) for font handling. Data handling and file input/output used data.table (v1.18.2.1), dplyr (v1.2.0) and the tidyverse (v2.0.0, RRID:SCR_019368), stringr (v1.6.0), openxlsx (v4.2.8.1) and yaml (v2.3.12). All analyses were performed in R (v4.5.2, RRID:SCR_001905).

The authors used Claude Code (Anthropic) as a coding assistant during development of the analysis scripts for this study. All code was specified, reviewed and tested by the authors, who verified that outputs were correct and reproducible. The authors take full responsibility for the analysis and for the content of this manuscript.

## RESULTS

### LOY in human ccRCC and across cancer entities is transcriptionally silent

To investigate the consequences of LOY in ccRCC, we leveraged bulk RNA-sequencing data from male tumors in the TCGA-KIRC [32] (n = 338) and CPTAC-ccRCC [33] (n = 137) cohorts. As previously described [13], we used the expression of nine stably expressed Y-chromosomal genes (*DDX3Y*, *UTY*, *KDM5D*, *USP9Y*, *ZFY*, *RPS4Y1*, *TMSB4Y*, *EIF1AY*, and *NLGN4Y*) to assess Y chromosome presence at the transcriptional level. The median scaled Y-chromosomal gene expression was bimodal in each cohort, allowing us to classify samples as loss (LOY) or retention (ROY) of the Y chromosome (**Additional file 1: Fig. S1A**). We excluded two outlier samples from the TCGA cohort (leaving n = 336) that had very low Y-chromosome expression scores, likely reflecting mislabeling or sex-discordance; one of them was independently flagged as sex-discordant [12]. For TCGA, the RNA-seq-based classification agreed with published WGS-based LOY calls for 95% of samples [12]. We have previously shown that *KDM5C* mutations are associated with LOY in ccRCC [34]. To avoid confounding by *KDM5C* status we excluded all samples harboring protein-altering somatic mutations in *KDM5C* or missing WES/WGS data from the analysis, retaining 289 TCGA-KIRC and 114 CPTAC-ccRCC tumors.

In both cohorts, we observed a similar rate of LOY (36.7% TCGA, 35.1% CPTAC), consistent with the previously reported frequency of about 40% [12]. The LOY group was significantly older (median 63 vs. 57 years, Wilcoxon p = 2 x 10^-4^). Tumor grade and stage were not associated with LOY status (**Additional file 1: Fig. S1D**). Unadjusted and age-adjusted overall survival did not differ between LOY and ROY (**Additional file 1: Fig. S1E, F**).

To test whether Y chromosome loss influences the transcriptome, we performed non-parametric differential expression analysis in each cohort (see Methods). We observed a strong relationship between the Y-gene score and tumor purity in the LOY but not in the ROY group, likely reflecting the dilution of the LOY signal by Y-carrying non-malignant cells (**Additional file 1: Fig. S1B**). To mitigate confounding by differences in infiltrating immune cells with potential concurrent Y-loss [13], we regressed out tumor purity and age effects prior to testing, applying the Wilcoxon rank-sum test to the resulting residuals. Per-cohort results for 16,176 protein-coding genes were then combined using Stouffer meta-analysis **(Additional file 2: Table S1)**. This revealed the expected strong downregulation of male-specific Y chromosomal (MSY) genes, including all nine used for Y-loss classification plus *TBL1Y* and *PRKY* (**Fig. 1A**). *PCDH11Y*, located at the X-transposed region of the Y chromosome, a region with 99% sequence similarity to Xq21 [35], was the only MSY gene not altered, possibly reflecting its low expression and mapping problems (**Additional file 1: Fig. S2A, B**). We next examined whether X-linked gametologs are upregulated in response to the loss of their Y counterparts. Only *DDX3X* (log2FC = 0.13, FDR = 0.017) and *TXLNG* (log2FC = 0.14, FDR = 0.011) showed significant upregulation in LOY tumors. Others, including *RPS4X*, *USP9X*, *KDM6A* (*UTX*), *ZFX*, *EIF1AX*, *PRKX*, and *TMSB4X*, showed no significant changes (**Additional file 1: Fig. S2C**).

**Figure 1.**
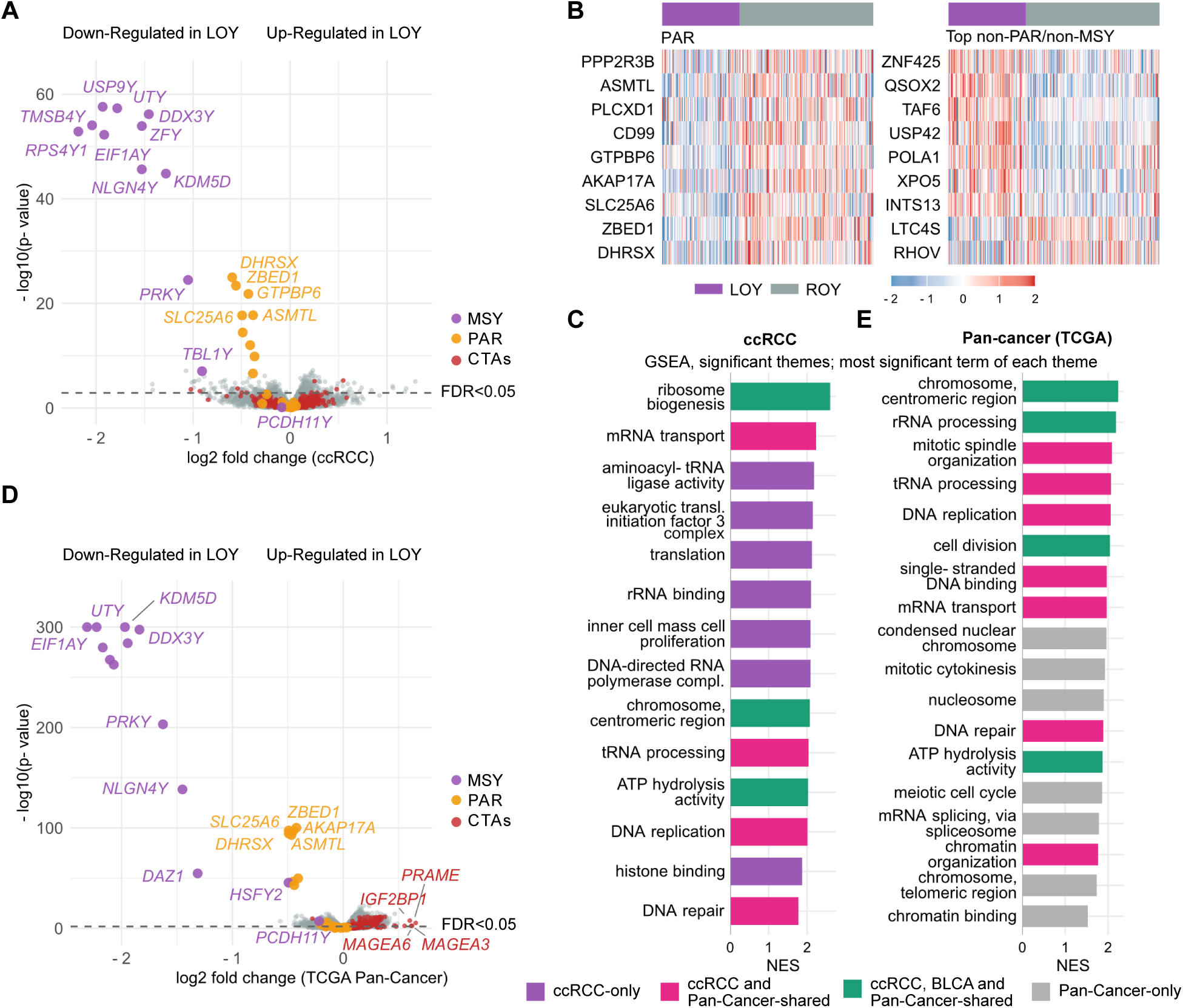
Loss of Y chromosome (LOY) in ccRCC and across cancer is not associated with consistent autosomal gene alterations that exceed pseudoautosomal dosage. **A** Volcano plot of the Stouffer meta-analyzed age- and purity-adjusted non-parametric differential expression analysis of LOY vs. ROY in male ccRCCs from TCGA KIRC and CPTAC ccRCC, restricted to *KDM5C*-wild-type. Benjamini-Hochberg adjusted false discovery rate (FDR) threshold of 0.05. **B** Z-score heatmaps of expression for nine FDR-significant pseudoautosomal region (PAR) genes (left panel) and of the same number of top-ranked non-PAR genes (right panel). **C** Normalized enrichment score (NES) of the most significant Gene Ontology (GO) term of each GO term theme of the GSEA. Positive scores indicate upregulation in LOY. Color-coded by overlap with significant bladder cancer (BLCA) and pan-cancer GO terms. **D** The same as A for Stouffer meta-analysis across TCGA pan-cancer entities, which include KIRC and BLCA. Genes that are testable in at least 90% of the cohorts (see Methods) were meta-analyzed. E The same as C for the significant pan-cancer GO term themes.

Among the non-MSY genes, the eight most significantly downregulated genes in LOY tumors (*DHRSX*, *ZBED1*, *ASMTL*, *GTPBP6*, *SLC25A6*, *AKAP17A*, *CD99*, *PPP2R3B*) were all located in the pseudoautosomal regions (PAR), which have autosomal characteristics and are present on both the X and Y chromosomes. This illustrates appropriate power of our analyses to detect mRNA changes associated with a halving of the gene dosage of PAR genes. From the 14 PAR1 genes, only three genes (*CRLF2*, *CSF2RA*, *P2RY8*) showed full dosage compensation in both datasets independently. None of the five PAR2 genes were altered upon LOY (**Additional file 1: Fig. S2D, E**). No autosomal gene reached the significance of the non-compensated PAR1 genes, and the top-ranked autosomal genes were directionally heterogeneous (**Fig. 1B**). We made similar observations in bladder cancer, where, in contrast to ccRCC, LOY is a strong prognostic marker [11] (**Additional file 1: Fig. S3A– F; Additional file 2: Table S2**).

Next, we applied gene set enrichment analysis (GSEA) to 4,242 Gene Ontology (GO) terms **(Additional file 2: Table S3)** to identify pathway alterations associated with LOY. In total, 35 GO terms reached Bonferroni-adjusted significance, all upregulated in LOY. The terms clustered into 14 functional themes including ribosome biogenesis, DNA repair, ATP hydrolysis activity, mRNA transport and cell division (**Fig. 1C, Additional file 1: Fig. S4A**). These pathway-level signals were, however, modest at the level of individual genes. The genes contributing most to these enrichments (the leading-edge genes) showed only small effect sizes (**Additional file 1: Fig. S4B**, median log_2_ fold change 0.1). The LOY-associated pathway response is therefore a coordinated but low-amplitude shift rather than a switch-like transcriptional program. The same analysis in TCGA-BLCA yielded 23 terms organized in eight themes (all upregulated in LOY), of which three themes (cell division, ribosome biogenesis, ATP hydrolysis activity) were shared with ccRCC (**Additional file 1: Fig. S4C, D; Additional file 2: Table S4**), defining a conserved pathway-level effect of LOY.

To test whether this generalizes to other cancer entities, we applied the same non-parametric differential expression analysis across male TCGA pan-cancer entities using WGS-based LOY calls [12] and combined them by Stouffer meta-analysis (**Additional file 2: Table S5)**. No autosomal gene exceeded an absolute log_2_FC of 1 (**Fig. 1D**). The most strongly altered, significant non-PAR/MSY genes were increased cancer/testis antigen transcripts (CTAs, e.g., *MAGEA6*, *MAGEA3, PRAME*), an association confined to stomach, liver, prostate, lung and bladder cancer and absent in ccRCC **(Additional file 1: Fig. S5A, B**). Gene set enrichment analysis (GSEA) identified 41 significant GO terms (all upregulated in LOY, **Additional file 2: Table S6**), converging on 18 themes including cell division, rRNA processing, DNA repair and ATP hydrolysis activity **(Fig. 1E, Additional file 1: Fig. S6A)**. Eleven pan-cancer themes overlapped with ccRCC themes. The absence of strong regulatory effects on individual genes, and the presence of low-amplitude cell cycle and chromosome segregation signals, are therefore not specific to ccRCC.

### LOY in an autochthonous murine ccRCC model is transcriptionally silent

A caveat of the conclusions from the transcriptomic analyses of human ccRCC is that genetic heterogeneity amongst ccRCC tumors might obscure the transcriptional consequences of LOY. We therefore investigated whether LOY also arises in the more genetically homogeneous background of ccRCC tumors from mice with a renal epithelium-specific deletion of *Vhl/Trp53/Rb1* [36]. The murine Y chromosome has substantial differences from the human Y chromosome, including about 700 protein-coding, mostly ampliconic genes and only one small, strain-specific pseudoautosomal region [37]. We defined stably expressed Y-chromosomal genes from RNA-seq of healthy murine kidney cortex (**Additional file 1: Fig. S7A**) and calculated a LOY score from *Ddx3y*, *Eif2s3y*, *Kdm5d* and *Uty* to set thresholds for lower- and higher-than-normal expression. Applying these thresholds in RNA-seq datasets of 35 male tumors from four different cohorts identified 14 with LOY and three with a higher-than-normal expression, which we excluded, leaving 18 ROY tumors for comparison (**Fig. 2A; Additional file 1: Fig. S7B; Additional file 2: Table S7**). The 40% frequency of LOY in the mouse ccRCC model is strikingly similar to the frequency observed in human ccRCC. PCA did not show clustering based on Y chromosome status (**Additional file 1: Fig. S7C**). Differential expression analysis of 13,446 genes revealed that, except for the four MSY genes, only 14 autosomal genes and one X-chromosomal gene (Msl3) were significantly altered (**Fig. 2B, Additional file 2: Table S8)** with considerable heterogeneity of expression amongst individual tumors (**Fig. 2C**). GSEA identified 19 significant terms (**Additional file 2: Table S9)** grouped into six themes including peptide antigen binding, cellular response to interferon beta and cellular response to exogenous dsRNA (all upregulated in LOY**, Additional file 1: Fig. S7D).** Notably, two of these themes, ribosome translation and chromosome segregation, are overlapping with human ccRCC, while all other terms were not even nominally significant in the human ccRCC analysis (**Fig. 2D**). Heatmaps of the leading-edge genes of these terms showed substantial expression heterogeneity within both the LOY and the ROY groups, with individual high- and low-expressing tumors distributed across both groups (**Additional file 1: Fig. S7E**).

**Figure 2.**
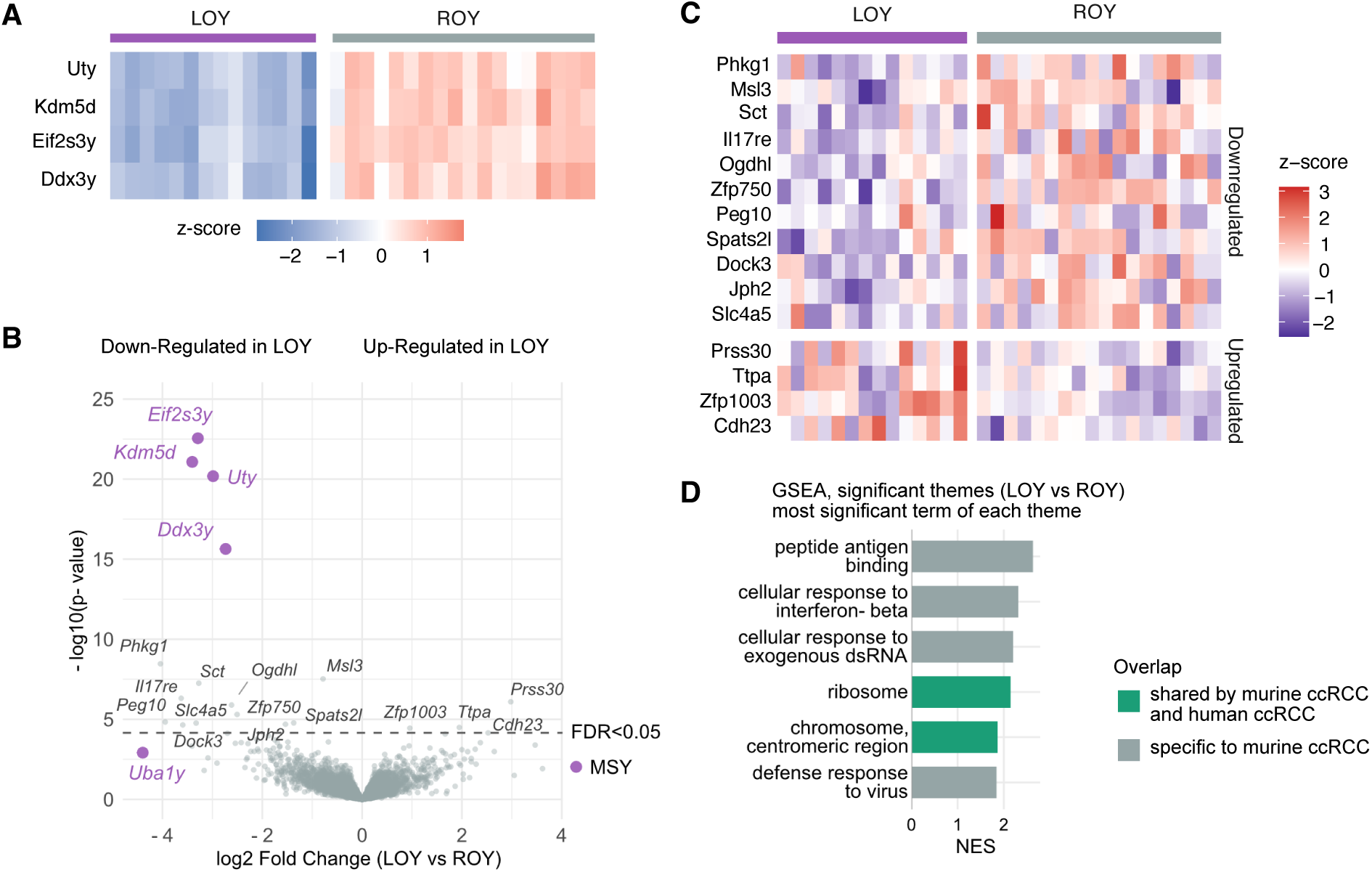
Spontaneous loss of Y (LOY) in autochthonous murine ccRCCs is transcriptionally silent. **A** Heatmap of row-wise Z-scored expression of four stably expressed MSY genes from RNA-seq of murine autochthonous ccRCCs. **B** Volcano plot of the differential expression analysis of LOY (n = 14) versus ROY (n = 18) tumors. Male-specific Y (MSY) genes are shown in purple; significantly altered autosomal genes (Benjamini-Hochberg adjusted) are named individually. **C** Heatmap of all differentially expressed autosomal genes (FDR < 0.05), row-wise Z-scored. **D** Normalized enrichment score of the most significant GO term of each GO term theme of the murine GSEA. The green color-coding indicates the overlap with significant human ccRCC terms.

### LOY in 786-O ccRCC cells is transcriptionally silent

Bulk RNA-seq of human and mouse ccRCC tumors could lack the resolution to detect cell-intrinsic transcriptional consequences of LOY, because of mRNA contributed by cells of the tumor microenvironment. To address this and to study LOY in an isogenic setting, we used CRISPR/Cas9 in an attempt to eliminate the Y chromosome in the human male ccRCC 786-O cell line. We cloned an sgRNA targeting 11 genes of the multi-copy TSPY gene family on the short arm of the Y chromosome (sgTSPY) into a lentiviral MuLE vector together with Cas9 and a neomycin resistance cassette (Fig. 3A). As controls, we used a MuLE vector carrying a non-targeting scramble sgRNA (sgCtrl) as well as inactive sgRNAs that lacked an adjacent PAM motif targeting the multi-copy RBMY1A1 gene family (sgRBMY1A1) or targeting the Y chromosomal centromere (sgY-Cen). To identify cells with LOY, we generated single cell clones and screened them by western blotting for KDM5D (Fig. 3B) and by RT-qPCR for RPS4Y1, DDX3Y and KDM5D (Fig. 3C, Additional file 2: Table S10). These genes lie on different arms of the Y chromosome (Fig. 3A). Screening identified five LOY clones across the four groups: 3/54 sgTSPY, 1/52 sgCtrl, 1/37 sgRBMY1A1 and 0/36 sgY-Cen (Fig. 3B, 3C). The isolation of LOY clones from the control-infected cells implies that 786-O cells can spontaneously lose the Y chromosome. Importantly, because LOY arose spontaneously in these clones, the Y-deficient clones analyzed here are not confounded by CRISPR/Cas9 cutting at the Y chromosome itself, and all comparisons are made between clones of the same parental line that differ in Y chromosome status. We conducted RNA-seq of the five clones with LOY and a selection of five ROY clones. We confirmed the Y chromosome status of all clones by applying the same Y score approach (**Additional file 1: Fig. S8A; Additional file 2: Table S11**). There was no clustering of the samples in the principal component analysis based on Y chromosome status **(Additional file 1: Fig. S8B).** Differential expression analysis revealed the expected downregulation of both MSY genes and seven of the nine non-compensated PAR1 genes identified in the human cohorts (except *PPP2R3B* and *PLCXD1*, which were the least expressed in ROY) (**Fig. 3D, Additional file 2: Table S12)**. Besides the Y chromosome-encoded genes, a total of 40 autosomal genes were significantly differentially expressed (FDR < 0.05) (**Fig. 3E**). Only two, *MMP13* and *EDN2*, reached an absolute log_2_ fold change above 2 and were not significantly altered in the human and murine ccRCCs. GSEA identified a single, significantly altered gene set, endoplasmic reticulum lumen (upregulated in LOY, **Additional file 2: Table S13**), which was altered neither in the human cohorts nor in murine ccRCCs, further supporting that LOY does not substantially affect expression of non-Y chromosome-encoded genes that can be detected in small datasets.

**Figure 3.**
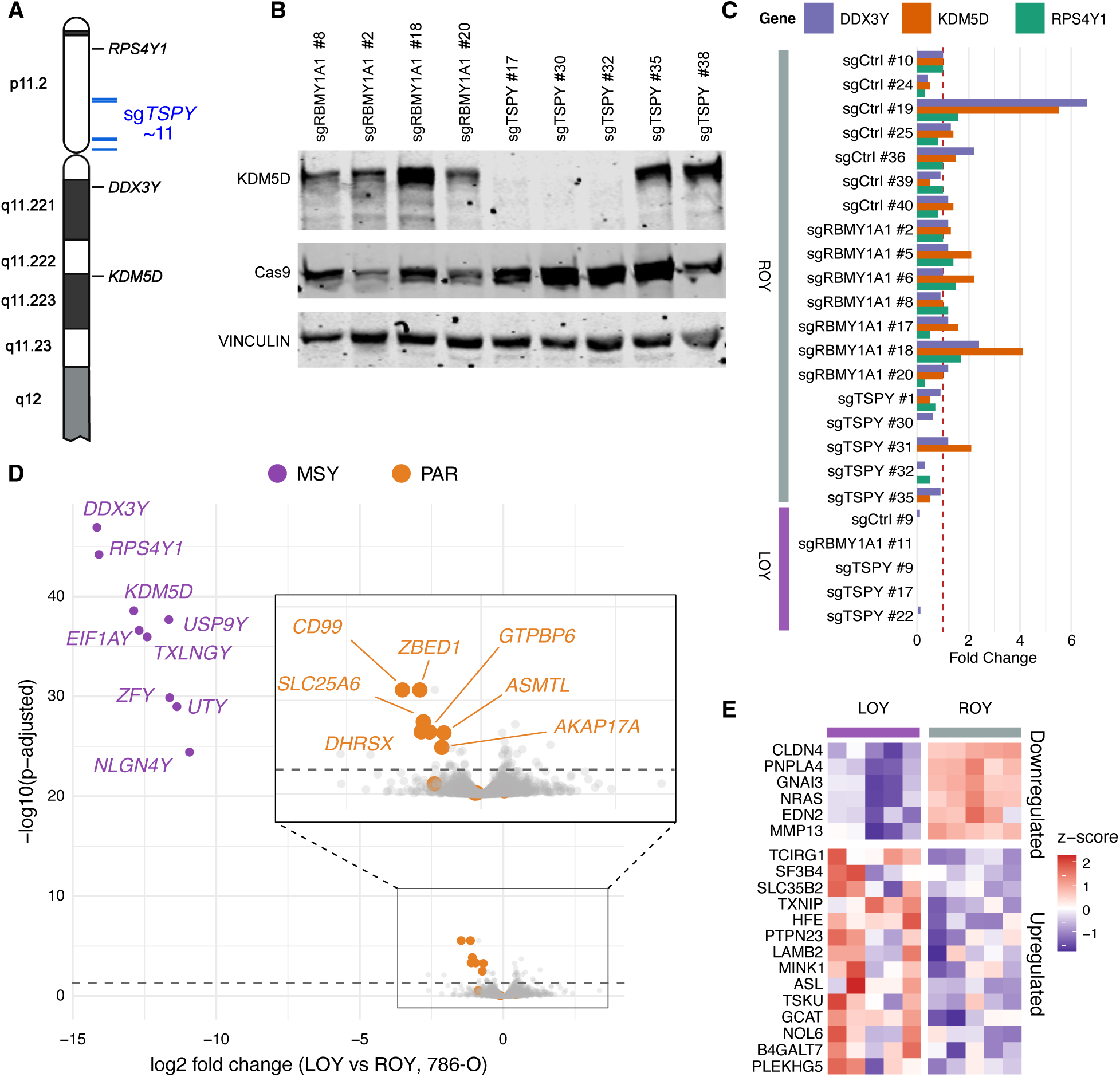
Loss of the Y chromosome in isogenic 786-O clones is transcriptionally silent. **A** Schematic of the location of the qPCR target genes (*RPS4Y1, DDX3Y, KDM5D*, in black) and sgRNAs (sgTSPY, in blue) along the regions of the Y chromosome. **B** Representative KDM5D and Cas9 western blots of a selection of screened single cell clones. **C** Fold change of expression of Y chromosome encoded *DDX3Y*, *KDM5D* and *RPS4Y1* upon normalization using *RPS18* and a control (sgCtrl #10). **D** Volcano plot of the differential expression analysis of LOY versus ROY cell lines. MSY genes are colored in purple, PAR genes in orange. Benjamini-Hochberg adjusted p-values. **E** Z-score heatmap of expression of the 20 most significantly, differentially expressed non-PAR non-MSY genes.

### LOY is associated with gametolog dependencies in ccRCC cells

Although LOY had little consequence for the transcriptome, the loss of Y-encoded gametologs could still create a dependency on their X-encoded counterparts. We therefore interrogated CRISPR gene-effect data from DepMap for the 11 expressed X-encoded gametologs across nine male ccRCC cell lines (3 LOY, 6 ROY; **Additional file 1: Fig. S9A–C**). *DDX3X*, *EIF1AX*, and *RPS4X* showed a stronger dependency in LOY lines (FDR < 0.05), a pattern reproduced for *EIF1AX* in 17 male bladder urothelial carcinoma lines (**Additional file 1: Fig. S9D, E**).

## DISCUSSION

The Y chromosome encodes several transcription factors and epigenetic regulators, giving rise to the hypothesis that LOY causes transcriptional alterations that contribute to cancer. We find, however, across human cohorts, murine tumors, and isogenic LOY clones, that LOY in ccRCC leaves the transcriptome largely unaffected, beyond the expected MSY and PAR dosage effects. That three complementary settings with orthogonal strengths (heterogeneous human tumors, a defined mouse model, and an isogenic cell system) converge on the same result makes it unlikely that our findings reflect lack of power or a single artifact. We identify a modest but reproducible pathway-level signature of LOY in human ccRCC tumors that includes cell division, ribosomal biogenesis and DNA repair genes and is shared across TCGA entities. Because its amplitude at the level of individual genes is small, we interpret it as a coordinated reaction rather than as evidence for a Y-chromosomal transcriptional program. Alternatively, these signatures might reflect pre-existing cellular states that enhanced the probability and tolerance of the initial LOY event, rather than being driven by LOY itself.

Interestingly, in some entities, most strongly in stomach adenocarcinoma, LOY was associated with a higher expression of cancer/testis antigens, which are normally silenced in somatic tissues and have previously been linked to LOY in lung adenocarcinoma [38]. This association, however, was absent in ccRCC, and its basis, possibly the global hypomethylation that accompanies aneuploidy, remains undetermined.

While potential cell type-specific or tumor entity-specific effects of LOY on transcription should not be ignored, we argue that future efforts to decipher the contribution of LOY to different tumor entities should have a particular focus on non-transcriptional functions of X- and Y-encoded homologs such as KDM5C/D and UTX/Y. Even though LOY might not be prognostically relevant in ccRCC, it still offers a potential vulnerability. In the ccRCC subgroup analysis of dependencies, we confirmed the previously described vulnerabilities to *EIF1AX*, *DDX3X and RPS4X* knockouts in ccRCC [39]. That the gametolog pair *RPS4Y1* and *RPS4X* encodes a structural component of the 40S ribosome is noteworthy in the context of our identification of upregulation of GSEA terms associated with ribosome biogenesis.

Our study has limitations. Y chromosome status in ccRCC was inferred from transcript abundance rather than DNA, so partial or subclonal losses are not resolved, although the 95% concordance with WGS-based calls argues that misclassification is limited. The human analyses rest on bulk RNA-seq. Despite purity adjustment, cell type-specific effects of LOY, particularly in infiltrating immune cells, would require single-cell data. However, the effect of LOY on immune cells is not subject to this study. Finally, the isogenic experiments used a single cell line, and all our readouts are transcriptional, so post-transcriptional effects of Y loss would go undetected.

## CONCLUSIONS

Our results reveal the unexpected finding that Y chromosomal genes are not prominently involved in the regulation of autosomal genes and motivate a shift in research focus toward other non-transcriptional consequences of LOY in cancer.

## List of Abbreviations

BLCA: bladder urothelial carcinoma (TCGA cohort)
ccRCC: clear cell renal cell carcinoma
cDNA: complementary DNA
CPTAC: Clinical Proteomic Tumor Analysis Consortium
CTA: cancer/testis antigen
Ct: cycle threshold
dsRNA: double-stranded RNA
ESTIMATE: Estimation of STromal and Immune cells in MAlignant Tumours using Expression data
FDR: false discovery rate
GDC: Genomic Data Commons
GEO: Gene Expression Omnibus
GO: Gene Ontology
GSEA: gene set enrichment analysis
KIRC: kidney renal clear cell carcinoma (TCGA cohort)
LOY: loss of the Y chromosome
log2FC: log2 fold change
MC3: Multi-Center Mutation Calling in Multiple Cancers
mLOY: mosaic loss of the Y chromosome
MSY: male-specific region of the Y chromosome
MuLE: multiple lentiviral expression (vector system)
NES: normalized enrichment score
PAM: protospacer adjacent motif
PAR: pseudoautosomal region
PCA: principal component analysis
qPCR: quantitative polymerase chain reaction
RNA-seq: RNA sequencing
ROY: retention of the Y chromosome
SD: standard deviation
sgRNA: single guide RNA
STAD: stomach adenocarcinoma (TCGA cohort)
STR: short tandem repeat
TCGA: The Cancer Genome Atlas
TPM: transcripts per million
WES: whole-exome sequencing
WGS: whole-genome sequencing
WT: wild-type

## Acknowledgements

We are grateful to the staff of the Lighthouse Core Facility for assistance with qPCR (funded in part by the Medical Faculty, University of Freiburg (Project Number 2025/B3-Fol)).

## Authors’ contributions

AW, PS and IJF conceived and designed the study. AW, RS, FC, PS and IJF developed the methodology. AW and AT performed the formal analysis of the RNA-seq data. RS and FC carried out the cell line experiments. ZRH developed the GSEA theme code. MA, LNZ and CW provided the murine tumor RNA-seq datasets. AW, RS and AT prepared the figures. AW, RS, AT, PS and IJF wrote the original draft. AW, RS, AT, FC, CW, ZRH, AK, MB, PS and IJF reviewed and edited the manuscript. FC, PS and IJF supervised the work. AK, MB, PS and IJF acquired funding. All authors read and approved the final manuscript.

Alex Waterhölter, Rebecca Scholz and Andreas Tsouris contributed equally to this work.

Correspondence to Pascal Schlosser or Ian J. Frew.

## Ethics declarations

### Ethics approval and consent to participate

TCGA/CPTAC are public de-identified data used under their data use policies, consent obtained by the consortia, no additional IRB approval required. No new animal experiments were performed for this study.

### Consent for publication

Not applicable.

### Availability of data and materials

Human data analyzed in this study are publicly available from the GDC data portal (https://portal.gdc.cancer.gov/). Data were used in accordance with the TCGA and CPTAC data use policies. DepMap data (release 25Q2) [40] is publicly available and was obtained from depmap.org. Murine raw RNA-seq datasets are publicly available under GSE344622, GSE330887, GSE150983 and [GEO ACCESSION TBD]. Raw RNA-seq data generated in this study comprising the 786-O clones have been deposited in the Gene Expression Omnibus under accession GSE346681. Analysis code is available at https://github.com/genepi-freiburg/Loss-of-the-Y-chromosome-is-transcriptionally-silent and archived at Zenodo.

### Authors’ information

Not applicable.

## Supplementary Information

**Additional file 1. Supplementary Figures S1 to S9 (PDF)**

Fig. S1 Y-loss classification and clinical association in male ccRCC. Fig. S2 Male-specific Y, pseudoautosomal gene and gametolog expression in ccRCC. Fig. S3 Differential expression analysis in TCGA bladder cancer. Fig. S4 GSEA themes in ccRCC and bladder cancer. Fig. S5 Expression of cancer/testis antigens in TCGA pan-cancer. Fig. S6 Pan-cancer GSEA themes. Fig. S7 LOY detection and its transcriptional impact in autochthonous murine ccRCC. Fig. S8 Y score and principal component analysis in 786-O clones. Fig. S9 DepMap screen for LOY-specific vulnerabilities.

**Additional file 2. Supplementary Tables S1 to S13 (XLSX)**

Table S1 Differential expression meta-analysis, human ccRCC (TCGA-KIRC and CPTAC-ccRCC, *KDM5C* wild-type, LOY vs ROY). Table S2 Differential expression, TCGA-BLCA (*KDM6A* wild-type). Table S3 GSEA, human ccRCC. Table S4 GSEA, TCGA-BLCA. Table S5 Differential expression meta-analysis across 27 male TCGA entities. Table S6 GSEA, TCGA pan-cancer meta-analysis. Table S7 Sample metadata and Y scores, murine RNA-seq. Table S8 Differential expression, murine VpR ccRCC (LOY vs ROY). Table S9 GSEA, murine ccRCC. Table S10 RT-qPCR fold changes of RPS4Y1, DDX3Y and KDM5D in 786-O clones. Table S11 Sample metadata and Y status, 786-O RNA-seq. Table S12 Differential expression, 786-O clones (LOY vs ROY). Table S13 GSEA, 786-O clones.

**Additional file 3. Uncropped western blots underlying Fig. 3B (PDF).**

