## Additional File 1 - Supplementary Figures for "Loss of the Y chromosome is transcriptionally silent in clear cell renal cell carcinoma and across cancer entities"

### Additional file 1: Supplementary Figures S1 to S9

**Contents**

1. **Fig. S1** Y-loss classification and clinical association in male ccRCC
2. **Fig. S2** Male-specific Y, pseudoautosomal gene and gametolog expression in ccRCC
3. **Fig. S3** Differential expression analysis in TCGA bladder cancer (BLCA)
4. **Fig. S4** GSEA themes in ccRCC and bladder cancer
5. **Fig. S5** Expression of cancer/testis antigens in TCGA pan-cancer
6. **Fig. S6** Pan-cancer GSEA themes
7. **Fig. S7** LOY detection and its transcriptional impact in autochthonous murine ccRCC
8. **Fig. S8** Y score and principal component analysis in 786-O
9. **Fig. S9** DepMap screen for LOY-specific vulnerabilities

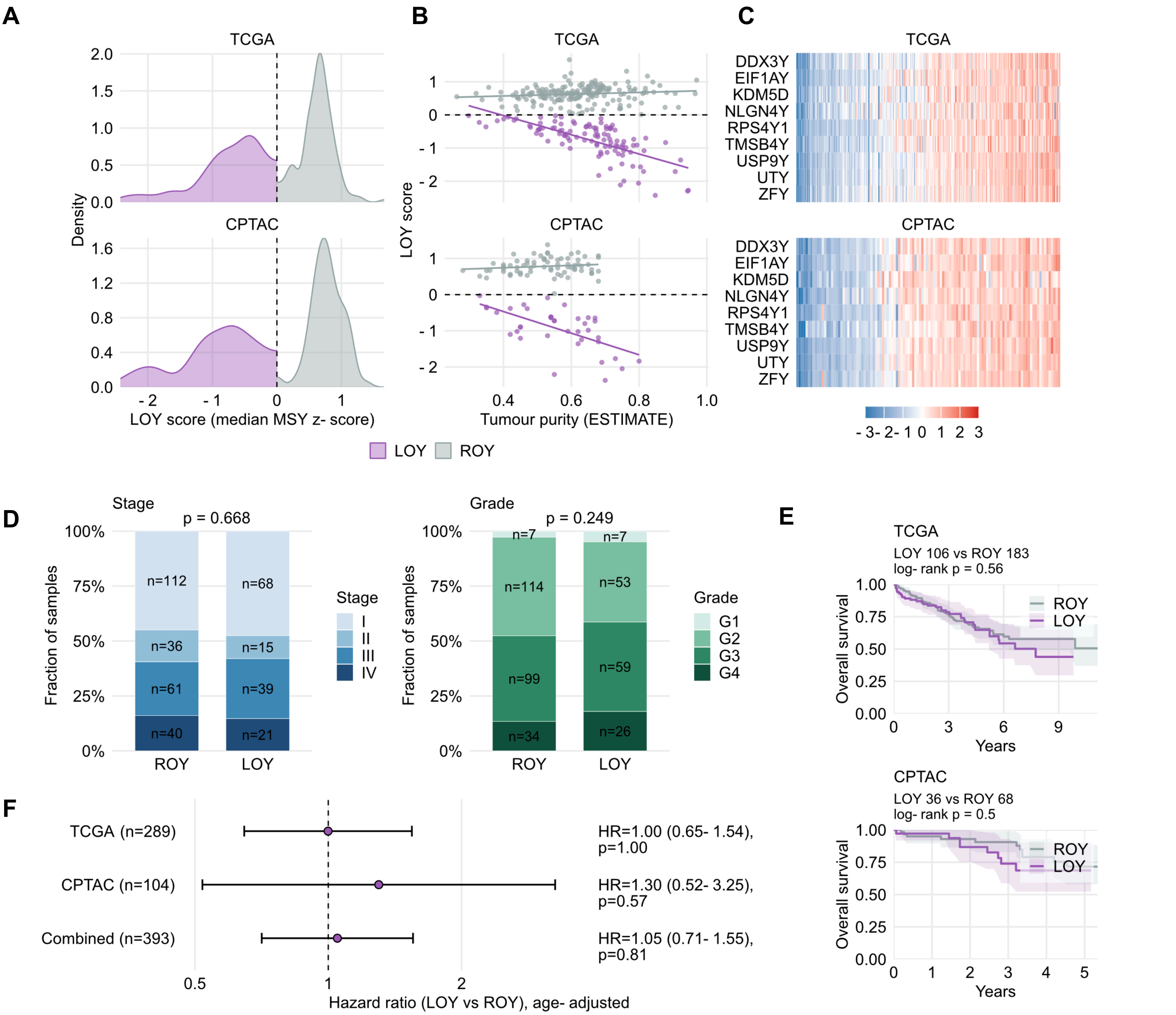


**Fig. S1 Y-loss classification and clinical association in male ccRCC.** All panels use KDM5C-wild-type (determined by WGS and/or WES; samples with missing data excluded) male ccRCC tumors from two cohorts: TCGA-KIRC (n = 289; 106 LOY, 183 ROY, after exclusion of 2 LOY score outliers with no detectable Y expression consistent with female sex) and CPTAC-ccRCC (n = 114; 40 LOY, 74 ROY). **A** Density of the continuous LOY score (median z-score of nine canonical MSY genes: *DDX3Y*, *EIF1AY*, *KDM5D*, *NLGN4Y*, *RPS4Y1*, *TMSB4Y*, *USP9Y*, *UTY*, *ZFY*). The vertical dashed line at score = 0 marks the LOY/ROY classification threshold. **B** LOY score versus ESTIMATE-derived tumor-purity per cohort, with separate linear fits for LOY (purple) and ROY (grey). Five CPTAC cases without a purity estimate were excluded. **C** z-scored expression heatmaps of the nine MSY scoring genes. Columns are sorted by LOY score (lowest = most LOY-like, leftmost). **D** Stacked proportional bar plots of tumor stage (n = 392) and grade (n = 399) by LOY status, pooled across TCGA-KIRC and CPTAC-ccRCC male patients, including patients with the respective variable recorded. Numbers inside each segment are sample counts within that LOY group; the annotation at the top of each panel is pooled, simulated Fisher's exact p-value (B = 50,000). **E** Kaplan–Meier overall-survival curves for LOY (purple) vs. ROY (grey) in male KDM5C-wild-type patients from TCGA-KIRC (top, n = 289) and CPTAC-ccRCC (bottom, n = 104). Restricted to patients with survival follow-up data. Shaded regions, 95% point-wise confidence intervals. Two-sided log-rank p shown in each panel subtitle. **F** Forest plot of age-adjusted Cox proportional-hazards estimates for LOY vs. ROY overall survival, fitted per cohort and combined across cohorts with stratification by cohort (n = 393). Points are hazard ratios with 95% confidence intervals.

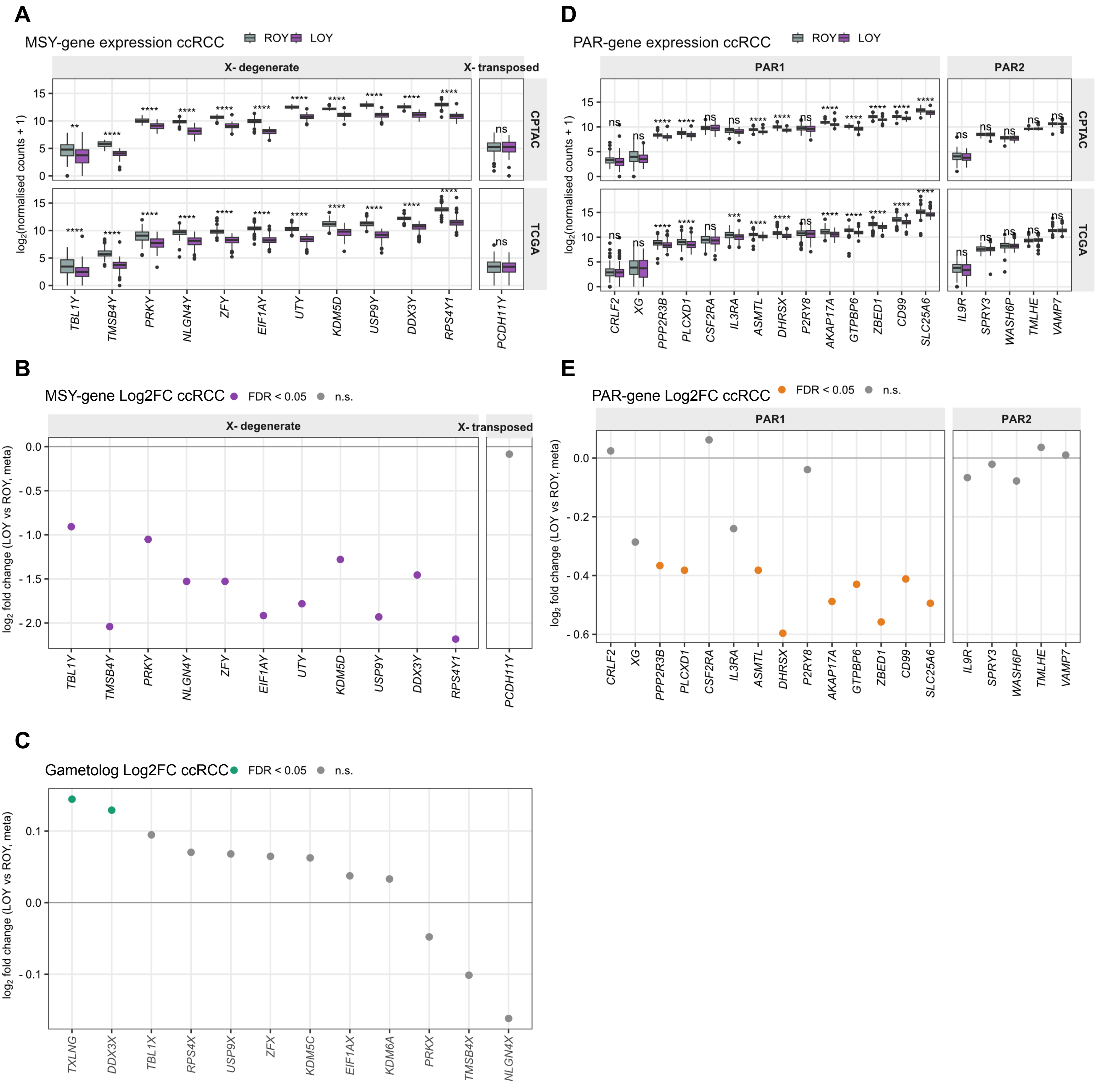


**Fig. S2 Male-specific Y, pseudoautosomal gene and gametolog expression in ccRCC.** All panels use KDM5C-wild-type tumors from male ccRCC patients with both differential-expression covariates (age at diagnosis, tumor purity) recorded (TCGA-KIRC n = 288, 106 LOY, 182 ROY; CPTAC-ccRCC n = 101, 36 LOY, 65 ROY). **A** Expression of MSY genes in LOY (purple) versus ROY (grey) tumors, as log2(normalized counts + 1), grouped by MSY sequence class. CPTAC-ccRCC, top; TCGA-KIRC, bottom. **B** Per-gene meta-analysis log2 fold change (LOY vs ROY) for MSY genes, ordered as in A. Purple, transcriptome-wide FDR < 0.05; grey, not significant. **C** Meta-analysis log2 fold change for the X-encoded gametologs. Green, FDR < 0.05 (transcriptome-wide BH). Twelve of the 16 canonical X-Y pairs are quantified in these cohorts; *AMELX*, *PCDH11X*, *TGIF2LX* and *VCX* are not. **D** As in A, for pseudoautosomal (PAR) genes, grouped into PAR1 and PAR2 (19 genes). **E** As in B, for PAR genes. Orange, FDR < 0.05. Boxplots in **A** and **D**: center line, median; box, IQR; whiskers, 1.5 × IQR; points, outliers. Asterisks denote two-sided Wilcoxon rank-sum tests, Benjamini-Hochberg-adjusted within cohort across the genes shown: ****$p < 10^{-4}$, ***$p < 10^{-3}$, **$p < 0.01$, *$p < 0.05$; ns, not significant.

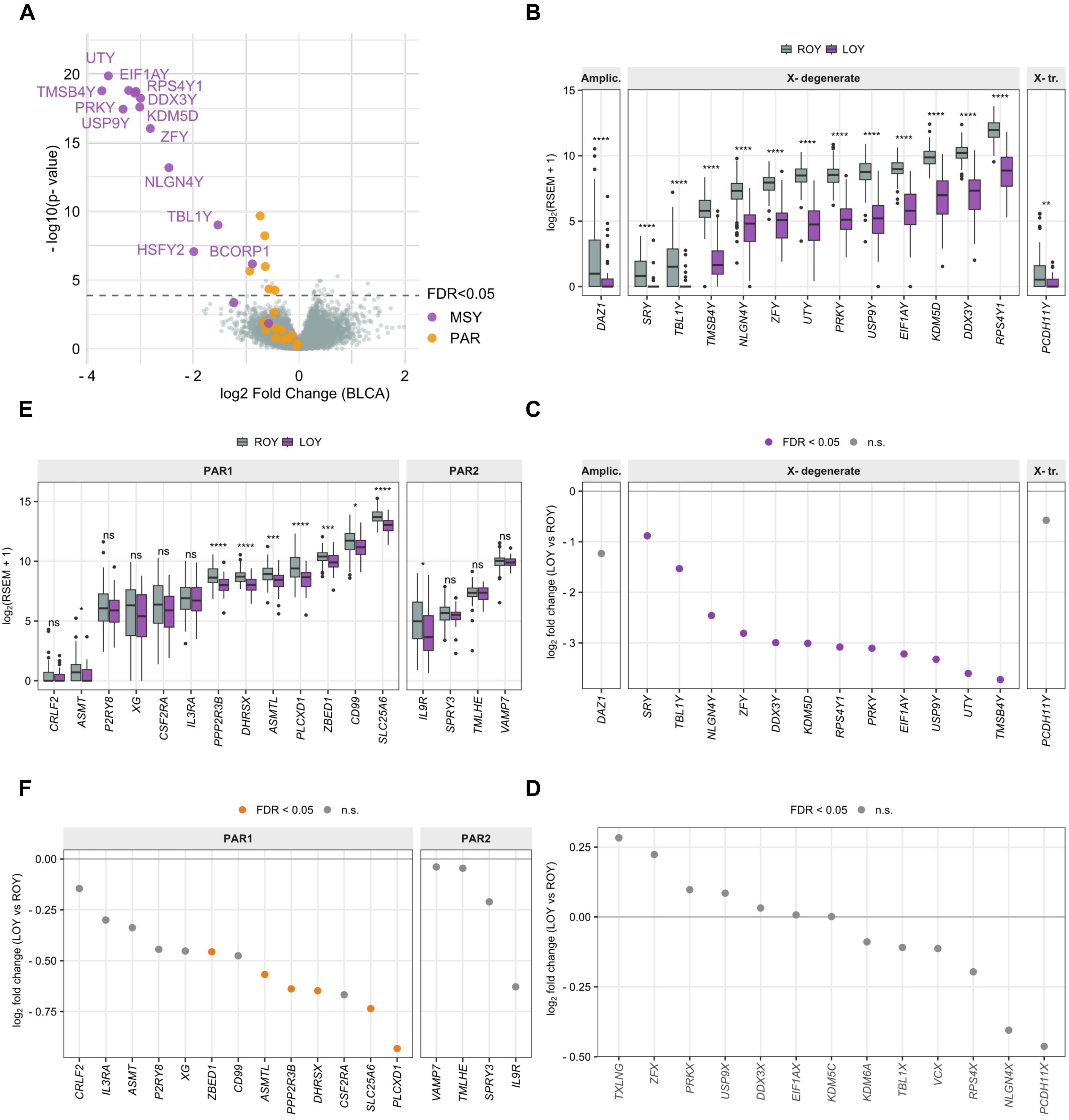


**Fig. S3 Differential expression analysis in TCGA bladder cancer (BLCA).** All panels use *KDM6A*-wild-type male TCGA-BLCA tumors (n = 167; 47 LOY, 120 ROY), with WGS-based LOY calls from Qi et al. **A** Volcano plot of LOY versus ROY differential expression (Wilcoxon rank-sum, age and tumor purity adjusted residuals). Purple, MSY genes; orange, PAR genes; grey, all other genes. Dashed line, FDR < 0.05. **B** Expression of male-specific region of the Y (MSY) genes in LOY (purple) versus ROY (grey) tumors, as $\log_2$(RSEM + 1), grouped by MSY sequence class - ampliconic, X-degenerate and X-transposed. **C** Per-gene $\log_2$ fold change (LOY vs ROY) for MSY genes, ordered as in B. Purple, FDR < 0.05. **D** $\mathrm{Log}_2$ fold change for the X-encoded gametologs; 13 of the 16 canonical X-Y pairs are quantified in BLCA (*TMSB4X*, *AMELX* and *TGIF2LX* are not). No gametolog reaches FDR < 0.05. **E** As in B, for pseudoautosomal (PAR) genes, grouped into PAR1 and PAR2. **F** As in C, for PAR genes. Orange, FDR < 0.05. Boxplots in B and E: Center line, median; box, IQR; whiskers, 1.5 × IQR. Asterisks denote two-sided Wilcoxon rank-sum tests, Benjamini-Hochberg-adjusted across the genes shown: ****$p < 10^{-4}$, ***$p < 10^{-3}$, **$p < 0.01$, *$p < 0.05$; ns, not significant.

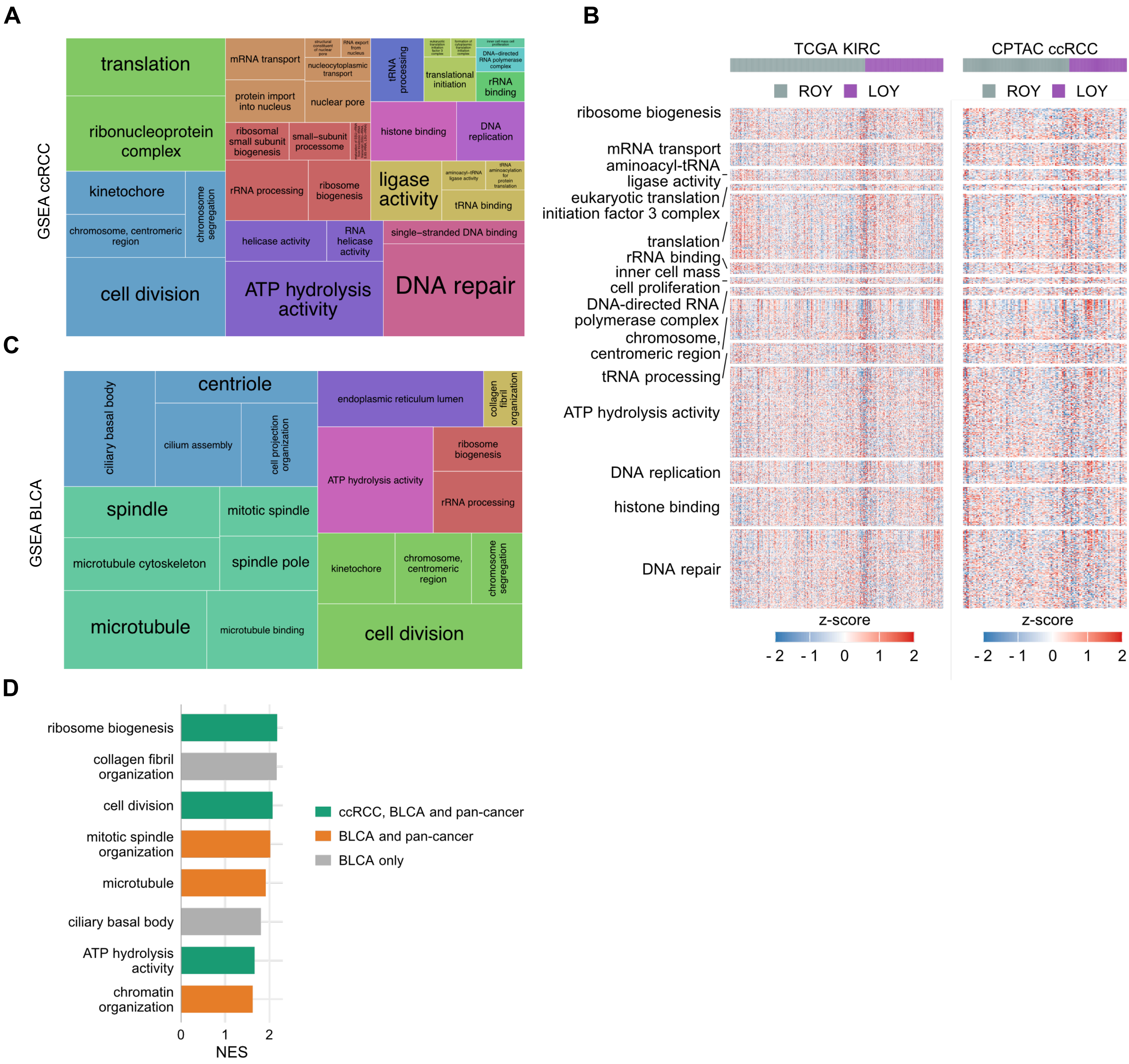


**Fig. S4 GSEA themes in ccRCC and bladder cancer. A** Treemap of the ccRCC LOY GSEA. Tile area is proportional to the number of genes annotated to the term; tiles are grouped and colored by theme. **B** Expression of the leading-edge genes of the 14 ccRCC themes in LOY (purple) versus ROY (grey) tumors, z-scored within cohort: TCGA-KIRC (*n* = 289, left) and CPTAC-ccRCC (*n* = 114, right), all KDM5C wild-type. Row blocks are labelled by most significant term of the theme; columns are ordered by LOY status. **C** As in A, for TCGA-BLCA. (d) Normalized enrichment scores (NES) of the BLCA themes, one bar per theme (represented by its name-giving term), colored by whether a member of the theme is also recovered in ccRCC, in all three analyses, or in BLCA alone.

A

B

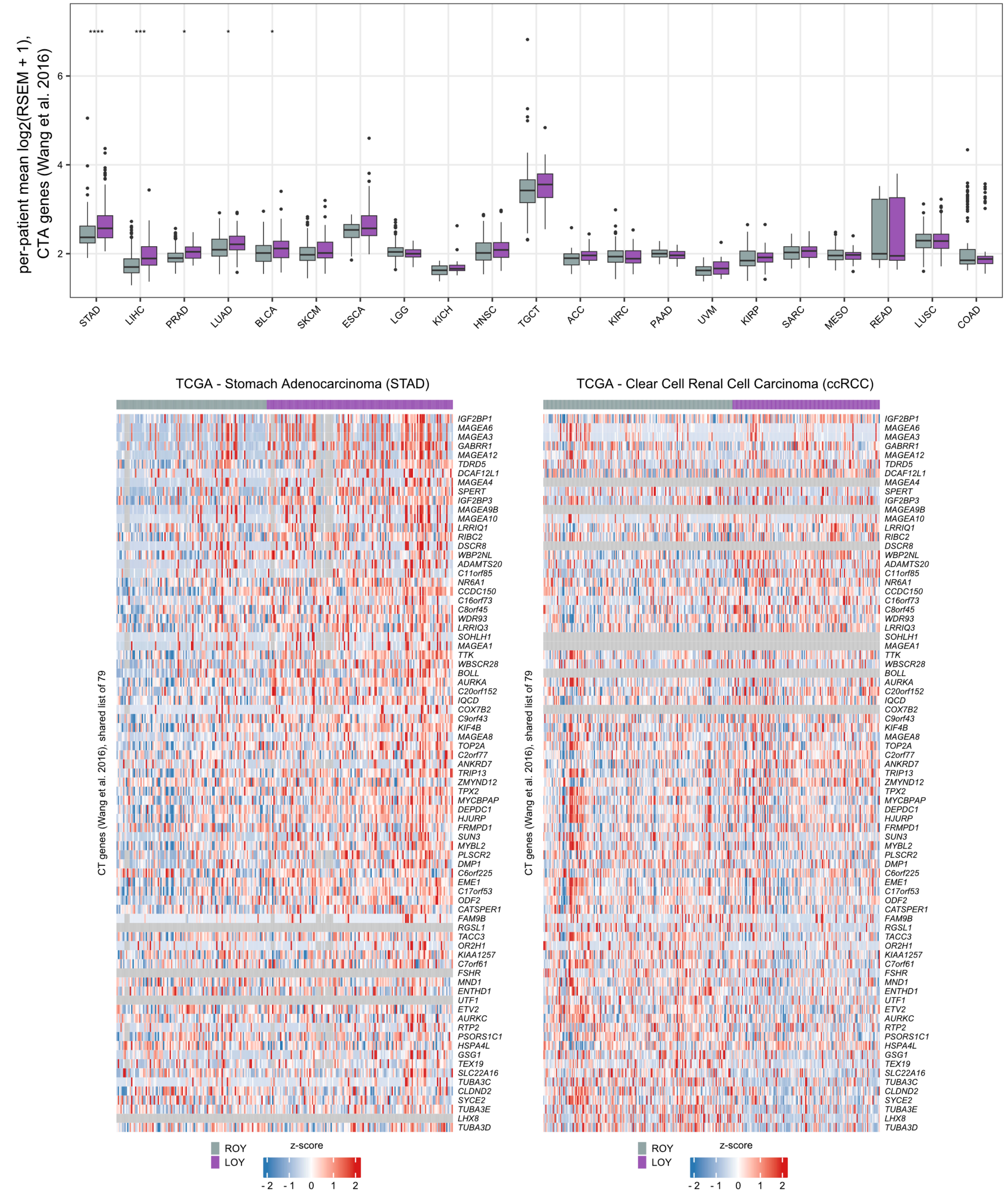


**Fig. S5 Expression of cancer/testis antigens in TCGA pan-cancer.** Cancer/Testis Antigen (CTA) genes are the set defined by Wang et al., Nat Commun 2016 (Supplementary Data 3A; 1,019 genes, of which 954 were quantified in the TCGA pan-cancer expression matrix). Eight of 29 entities were excluded for having fewer than 10 tumors in a group (BRCA, CHOL, DLBC, GBM, LAML, PCPG, THCA, THYM). **A** Per-patient CTA-module expression in LOY (purple) versus ROY (grey) tumors across 21 TCGA entities. Boxes show the median and IQR, whiskers extend to 1.5 × IQR, and points are outliers. Entities are ordered by significance. Asterisks denote two-sided Wilcoxon rank-sum tests on the per-patient scores, Benjamini-Hochberg-adjusted across the 21 entities (**** $p < 10{-}4$; *** $p < 10{-}3$; ** $p < 0.01$; * $p < 0.05$). **B** Expression of individual CTA genes in LOY versus ROY tumors in stomach adenocarcinoma (TCGA-STAD, n = 253; 140 LOY / 113 ROY) and clear cell renal cell carcinoma (TCGA-KIRC, n = 320; 140 LOY / 180 ROY), z-scored per gene within entity. Grey cells mark genes that are not expressed in that entity (log2 expression ≥ 1 in fewer than 10% of its samples), missing, or without variance. Y-linked genes are excluded.

**A**

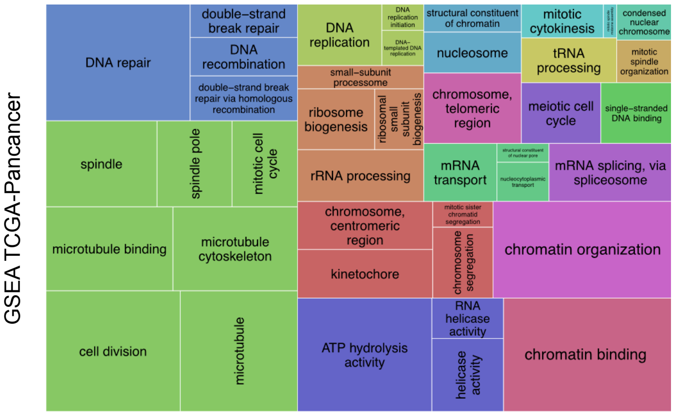


**Fig. S6 Pan-Cancer GSEA themes. A** Treemap as in Fig. S4A, for the TCGA pan-cancer LOY meta-analysis.

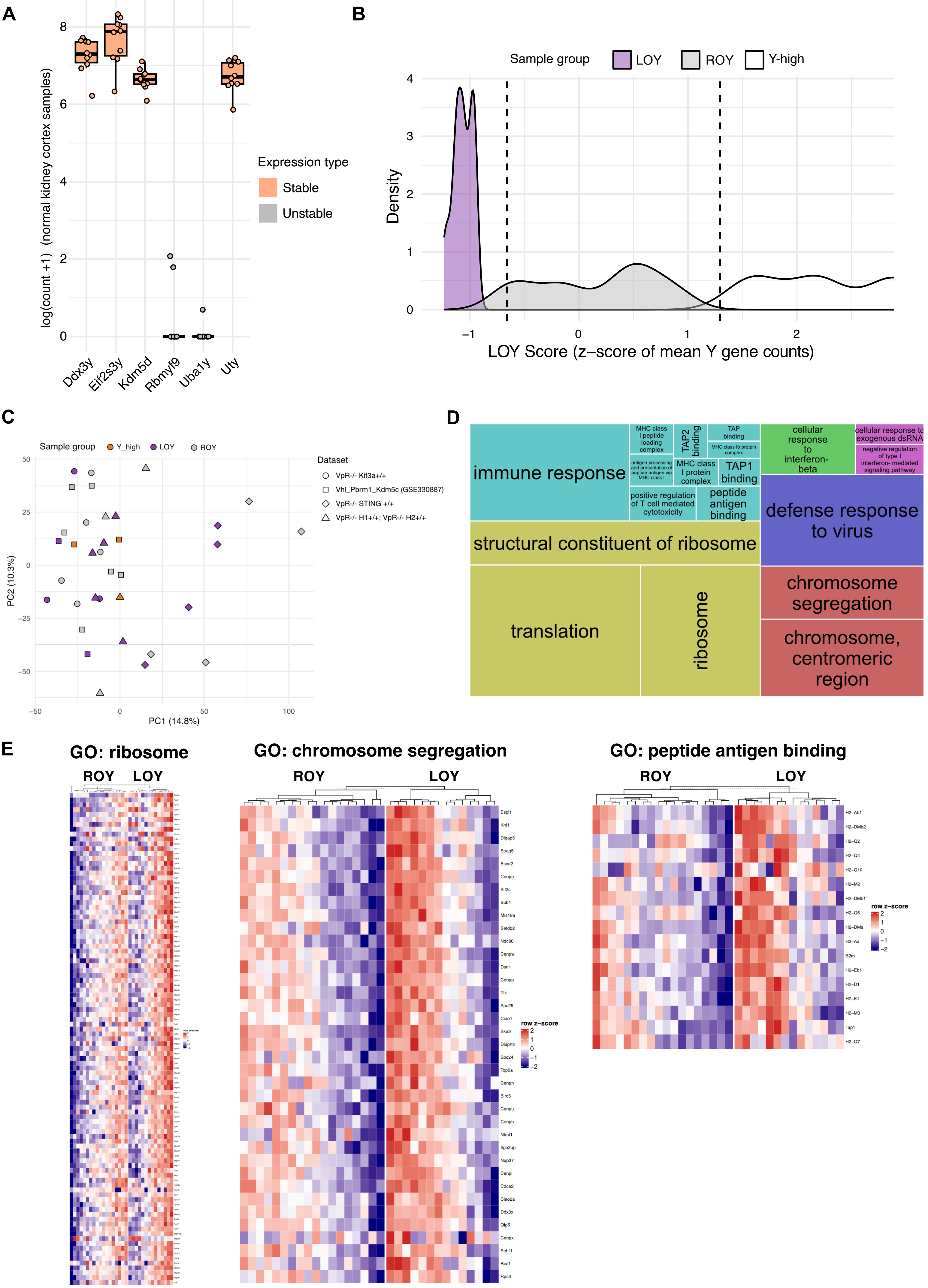
A
log(count +1) (normal kidney cortex samples)
8
6
4
2
0
Ddx3y
Eif2s3y
Kdm5d
Rbmyf9
Uba1y
Uty
Expression type
Stable
Unstable
B
Sample group
LOY
ROY
Y-high
Density
LOY Score (z−score of mean Y gene counts)
C
Sample group
Y_high
LOY
ROY
Dataset
VpR-/- Kif3a+/+
Vhl_Pbrm1_Kdm5c (GSE330887)
VpR-/- STING +/+
VpR-/- H1+/+; VpR-/- H2+/+
PC2 (10.3%)
PC1 (14.8%)
D
immune response
MHC class I peptide loading complex
TAP2 binding
TAP binding
MHC class Ib protein complex
antigen processing and presentation of peptide antigen via MHC class I
MHC class I protein complex
TAP1 binding
positive regulation of T cell mediated cytotoxicity
peptide antigen binding
cellular response to interferon-beta
cellular response to exogenous dsRNA
negative regulation of type I interferon- mediated signaling pathway
defense response to virus
structural constituent of ribosome
translation
ribosome
chromosome segregation
chromosome, centromeric region
E
GO: ribosome
ROY
LOY
GO: chromosome segregation
ROY
LOY
Espl1
Knl1
Dlgap5
Spag5
Esco2
Cenpo
Kif2c
Bub1
Mis18a
Setdb2
Ndc80
Cenpw
Dsn1
Cenpp
Ttk
Spc25
Ciao1
Ska3
Diaph3
Spc24
Top2a
Cenpn
Birc5
Cenpu
Cenph
Ntmt1
Itgb3bp
Nup37
Cenpi
Cdca2
Ciao2a
Ddx3x
Oip5
Cenpx
Seh1l
Rcc1
Rps3
row z−score
GO: peptide antigen binding
ROY
LOY
H2−Ab1
H2−DMb2
H2−Q2
H2−Q4
H2−Q10
H2−M2
H2−DMb1
H2−Q6
H2−DMa
H2−Aa
B2m
H2−Eb1
H2−D1
H2−K1
H2−M3
Tap1
H2−Q7
row z−score

**Fig. S7 LOY detection and its transcriptional impact in autochthonous murine ccRCC. A** Log-transformed expression of the MSY genes in RNA-seq of healthy murine kidney cortex. Genes showing stable expression (minimum count > 10) are in orange. **B** Categorization of the samples in 3 groups (LOY, ROY, Y-high) according to their LOY score. **C** Principal Component Analysis (PCA) plot of the murine samples. All genes in the Y chromosome were excluded in the calculation of the principal components. Symbols denote the originating experiment. **D** Treemap of the GO terms enriched in LOY versus ROY samples (19 Bonferroni-significant terms in six themes, clustered as in Supplementary Fig. 4A). **E** Heatmaps of the gene-wise z-scored expression of the leading-edge genes in the "Ribosome", "Chromosome Segregation" and "peptide antigen binding" gene sets.

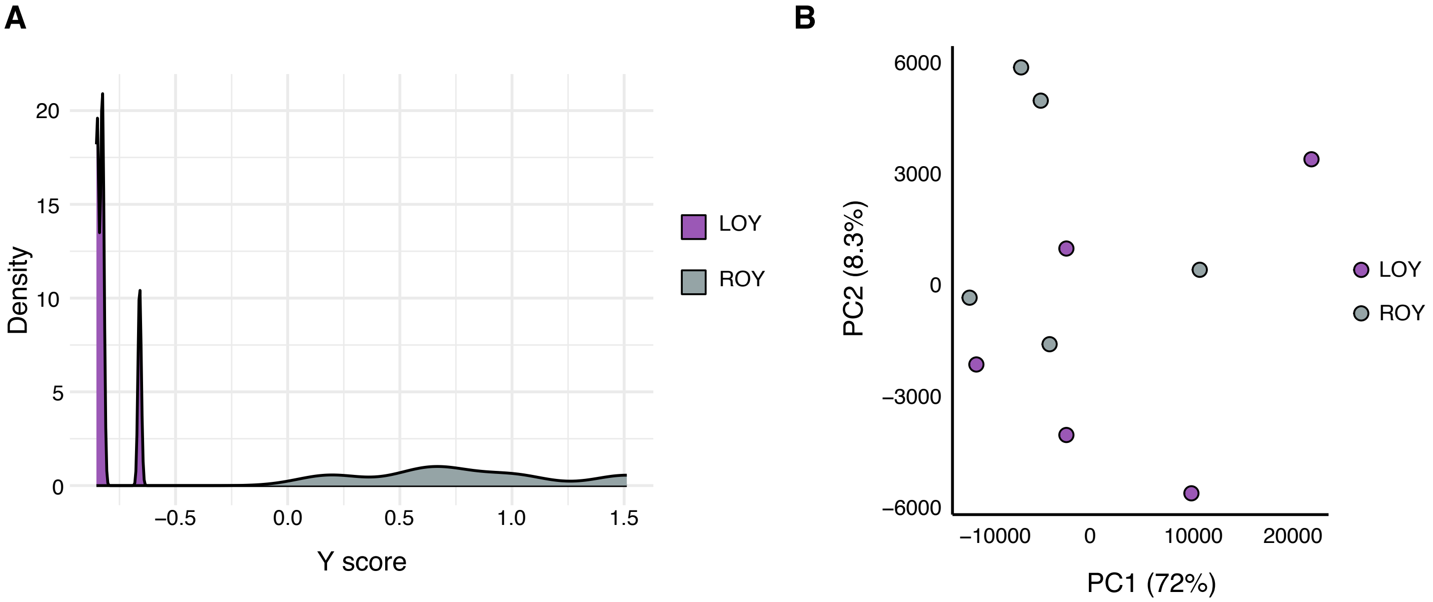


**Fig. S8 Y score and Principal Component Analysis in 786-O. A** Density plot of the Y scores. **B** Principal Component Analysis of the LOY and ROY samples. All Y chromosome genes were excluded in the calculation of the principal components.

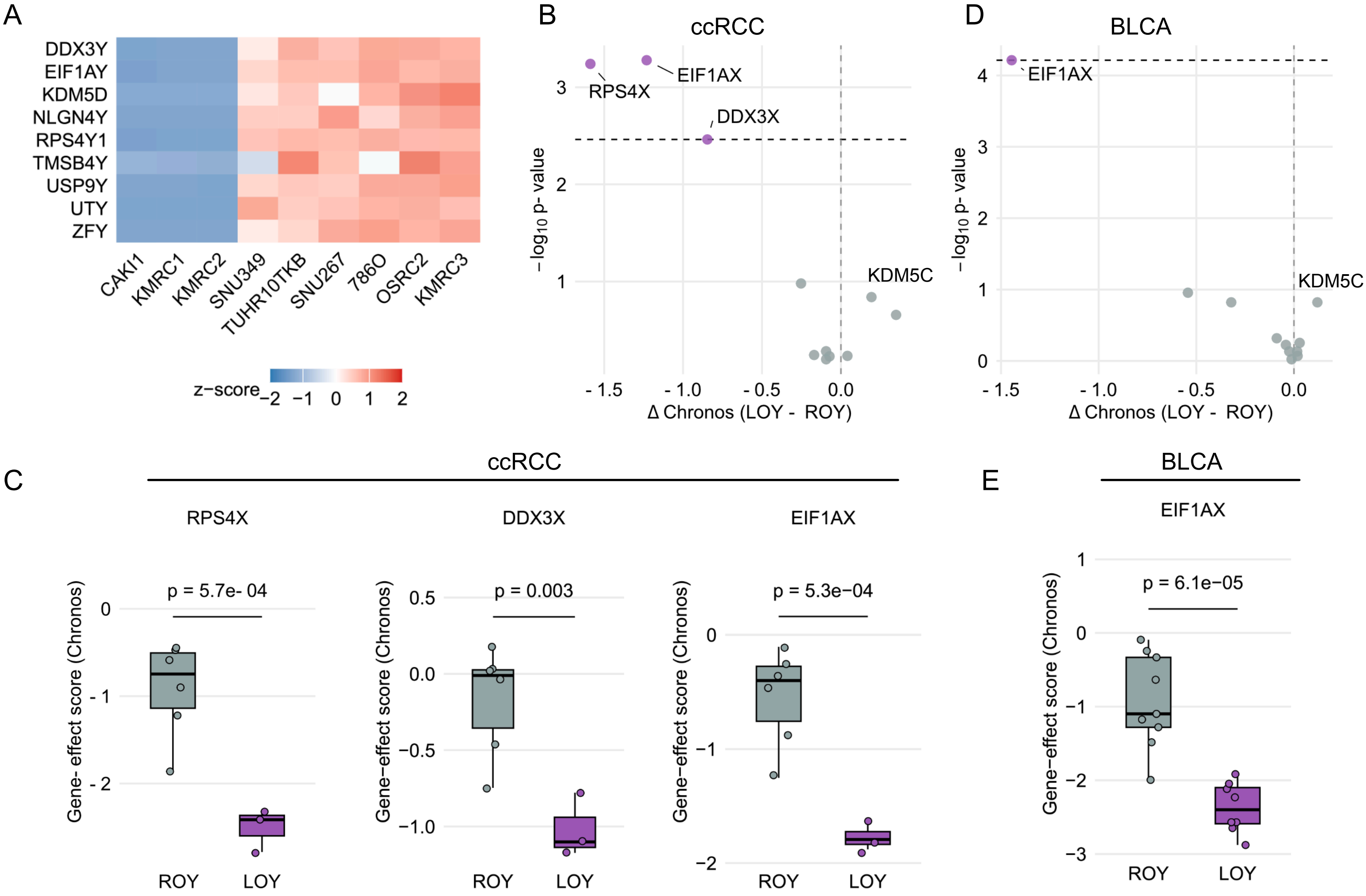


**Fig. S9 DepMap screen for LOY-specific vulnerabilities**. Analyses use CRISPR gene-effect (Chronos) scores from DepMap release 25Q2. The LOY score is the median z-score of nine canonical MSY genes (DDX3Y, EIF1AY, KDM5D, NLGN4Y, RPS4Y1, TMSB4Y, USP9Y, UTY, ZFY). **A** z-scored log2(TPM + 1) expression of the nine MSY scoring genes in nine male ccRCC cell lines. **B** Volcano plot of a gametolog-restricted, two-sided Welch's t-test on Chronos scores in those nine ccRCC lines (3 LOY vs 6 ROY). Testing was restricted to the 11 of 16 canonical X-Y gametolog pairs whose X copy is expressed in this cohort (log2(TPM + 1) ≥ 1 in ≥ 50% of lines): DDX3X, EIF1AX, KDM5C, KDM6A, PRKX, RPS4X, TBL1X, TMSB4X, TXLNG, USP9X and ZFX. Benjamini-Hochberg FDR was computed across these 11 genes; the dashed line marks FDR = 0.05. **C** Chronos scores in LOY (purple) versus ROY (grey) ccRCC lines for the three gametologs reaching FDR < 0.05: RPS4X, DDX3X and EIF1AX. Boxes show median and IQR, whiskers extend to 1.5 × IQR. **D** As in B, for 17 male bladder urothelial carcinoma (BLCA) cell lines (8 LOY vs 9 ROY). **E** As in C, for EIF1AX.
